# Rapid connectivity alterations of thalamic nuclei during initial learning of goal-directed behaviour

**DOI:** 10.64898/2026.05.15.725154

**Authors:** Chelsea Jarrett, Sofia Fregni, Katharina von Kriegstein, Hannes Ruge

## Abstract

The thalamus is essential for learning, dynamically engaging with other subcortical and cerebral cortex regions throughout the learning process. Here, the thalamus serves as a critical connector hub and synchroniser within the thalamocortical system of the brain. However, whilst higher order thalamic nuclei are known to be particularly important for this process, the exact contributions of individual higher order and first order thalamic nuclei, alongside their individual involvement with cortical networks and subcortical regions, remains unexplored within the initial phase of learning. In light of this, we analysed fMRI data obtained within a paradigm which is designed to examine initial learning processes within feedback-driven stimulus-response learning, in order to explore thalamic contributions. We investigated dynamic learning-related functional connectivity alterations between various thalamic nuclei with other subcortical regions and cortical networks. Our results show that the initial phase of learning was associated with: (1) decreasing functional connectivity between thalamic nuclei and frontoparietal and cingulo-opercular networks, (2) increasing functional connectivity between thalamic nuclei with default mode and salience networks, (3) decreasing functional connectivity between thalamic nuclei and the putamen, and (4) decreasing functional connectivity amongst higher order thalamic nuclei. Furthermore (5) these dynamic alterations were associated primarily by mediodorsal thalamus. Altogether, these results indicate that higher order thalamic nuclei play a crucial role within initial learning and in the generation of novel goal-directed behaviour. This was demonstrated through enhanced functional connectivity with selected cortical networks which drive goal-directed behaviour, alongside decreased functional connectivity with striatal regions which drive motor selectivity.

## Introduction

Goal-directed behaviour is voluntary action that is selected to achieve a desired goal (de Wit & Dickinson, 2009). This behaviour is often contingent upon the learning of associations between stimuli, responses and their outcomes in a certain context (Buschman & Miller, 2014; Burton *et al*., 2024), is driven by anticipated response outcomes, and is essential for flexible adaptation to dynamic environments (Griffiths *et al*., 2014).

During the initial phase of trial-and-error learning, a number of task processes are initiated. Namely, a period of exploration, whereby different motor responses must be tried out in order to derive correct versus error-related feedback. This therefore requires subjects to remember their responses and the outcomes of those responses (Cochrane & Green, 2023; Hassanzadeh *et al*., 2024) and places a high strain on working memory resources (Cowan, 2010; Ruge *et al*., 2018). However, with progressing exploration, identification of correct stimulus-response-outcome associations occur, and with their reinforcement, this strain on working memory decreases (Collins *et al*., 2017; Mohr *et al*., 2018). Across this initial learning phase, a dynamic process of downregulation and upregulation of exploratory response options is therefore occurring. First, at primary exposure to a specific stimulus, initially unchosen response options must be suppressed in favour of a single chosen response. Subsequently, previously chosen responses with unwanted outcomes have to be downregulated until the response yielding a desired outcome is eventually identified and subsequently further upregulated. In this paper, we specifically aim to describe the neural alterations that are associated with these kinds of initial learning processes, with a special focus on thalamic contributions.

In the thalamocortical system, the thalamus is connected to every region of the cerebral cortex (Hwang *et al*., 2017). In addition, the thalamus is also well-connected to subcortical systems, including the basal ganglia (Minagar *et al*., 2013). From an integrative network perspective, learning requires reciprocal flow of information from the thalamus to cortical and subcortical areas of the brain and is facilitated by complex feedback circuits called cortico-striato-thalamo-cortical (CSTC) loops (Baladron *et al*., 2020). These loops contain nodes which overlap with cortical networks (Raichle, 2015; Seeley, 2019; Butler *et al*., 2023), such as the salience network and its CSTC loop (Peters *et al*., 2016). In this way, these loops facilitate goal-directed behaviour by enabling global integration of information for flexible cognitive responses (Parkes *et al*., 2019). Furthermore, the thalamus is also supported in this special role as an integrative and modulatory hub for the rest of the brain (Saalmann, 2014) through its composition as a structure that is formed of numerous heterogeneous nuclei; each endowed with unique structural, functional (Fama & Sullivan, 2015; Zhang & Li, 2017) and connectivity properties (Hwang *et al*., 2017). Already, many of these nuclei have been identified as distinct nodes within various CSTC loops and cortical networks themselves (Kumar *et al*., 2022), and the diverse functional connectivity that the thalamus and its nuclei share with cortical and subcortical structures enables it to pull behaviour toward goal-directed or automatic control (La Terra *et al*., 2022). Accordingly, through its dynamic connections with cortical and subcortical regions, the thalamus is in an optimal position to drive initial learning processes and novel goal-directed behaviour through its reconfiguration of cortical networks (Kawabata *et al*., 2021). However, the specific contributions of individual thalamic nuclei and their associated functional connectivity alterations to these processes remain unexplored.

In the present paper, we analysed data from a recent study by Fregni *et al*. (2025), as this study utilised a learning paradigm that is designed to track the acquisition of novel goal-directed responses within the initial phase of learning, and is therefore suited to investigate potential functional connectivity alterations between thalamic nuclei, non-thalamic subcortical regions, and cortical networks. Here, we hypothesised that the initial phase of stimulus-response (S-R) learning would be associated with functionally diverse connectivity alterations between thalamic nuclei, non-thalamic regions, and cortical networks which are known to implement goal-directed control processes (Redgrave *et al*., 2010; Spreng *et al*., 2010; Fresno *et al*., 2019; Hausman *et al*., 2022; Crivelli-Decker *et al*., 2023). In particular, we expected to find increasing functional connectivity amongst higher order thalamic nuclei, basal ganglia, the hippocampus, and task-positive cortical networks. Finally, we also expected to find altered functional connectivity between thalamic nuclei and cortical regions with functionally-diverse subregions of the putamen.

## Materials and Methods

### Subjects

The sample consisted of a total of 78 participants (52 females, 26 males; mean age 23.97 years, SD = 4.42, range 18-34 years). Four additional subjects could not be used due to technical error or due to the participant terminating the session. All participants were right-handed, neurologically healthy, had normal or corrected vision, and had given their written consent for participation. The experimental protocol was approved by the Ethics Committee of the Technische Universität Dresden [EK586122019], and informed consent was obtained in writing from all subjects prior to their participation in the experiment. In addition, all participants were compensated 10 euros per hour, in addition to money that they had the opportunity to earn through their individual performance in the experiment.

### Experimental Procedure

The paradigm from Fregni *et al*. (2025) consisted of a rapid learning task, in which participants learned novel stimulus-response (S-R) associations within a short time-frame of just a few trials (8 repetitions per stimulus). The paradigm was split into three learning conditions, which included trial-and-error, observation-based, and instruction-based learning. However, our current study exclusively analysed the data obtained within the trial-and-error learning condition. For a detailed description of the observation based and the instruction-based learning conditions please refer to Fregni *et al*. (2025). Our focus solely on trial-and-error learning enabled comparison with results from a previous study which had also utilised a trial-and-error learning paradigm to examine thalamus-related connectivity alterations on a slower timescale of >80 repetitions per stimulus (Jarrett *et al*., 2025).

A total of 108 visual stimuli were used to create unique subject-specific randomized lists. These stimuli were black and white images of various animate or inanimate objects (Figure 1). They were distributed across 27 experimental blocks (including three practice blocks). Therefore, the full experiment comprised 24 blocks in total, 8 of which were trial-and-error blocks randomly intermixed with blocks of the other two learning conditions. The 24 blocks were presented across four functional runs while participants were lying inside the MR machine. Each functional run lasted 20 minutes and contained six blocks, formed of two blocks of each of the three learning conditions.

**Figure 1.**
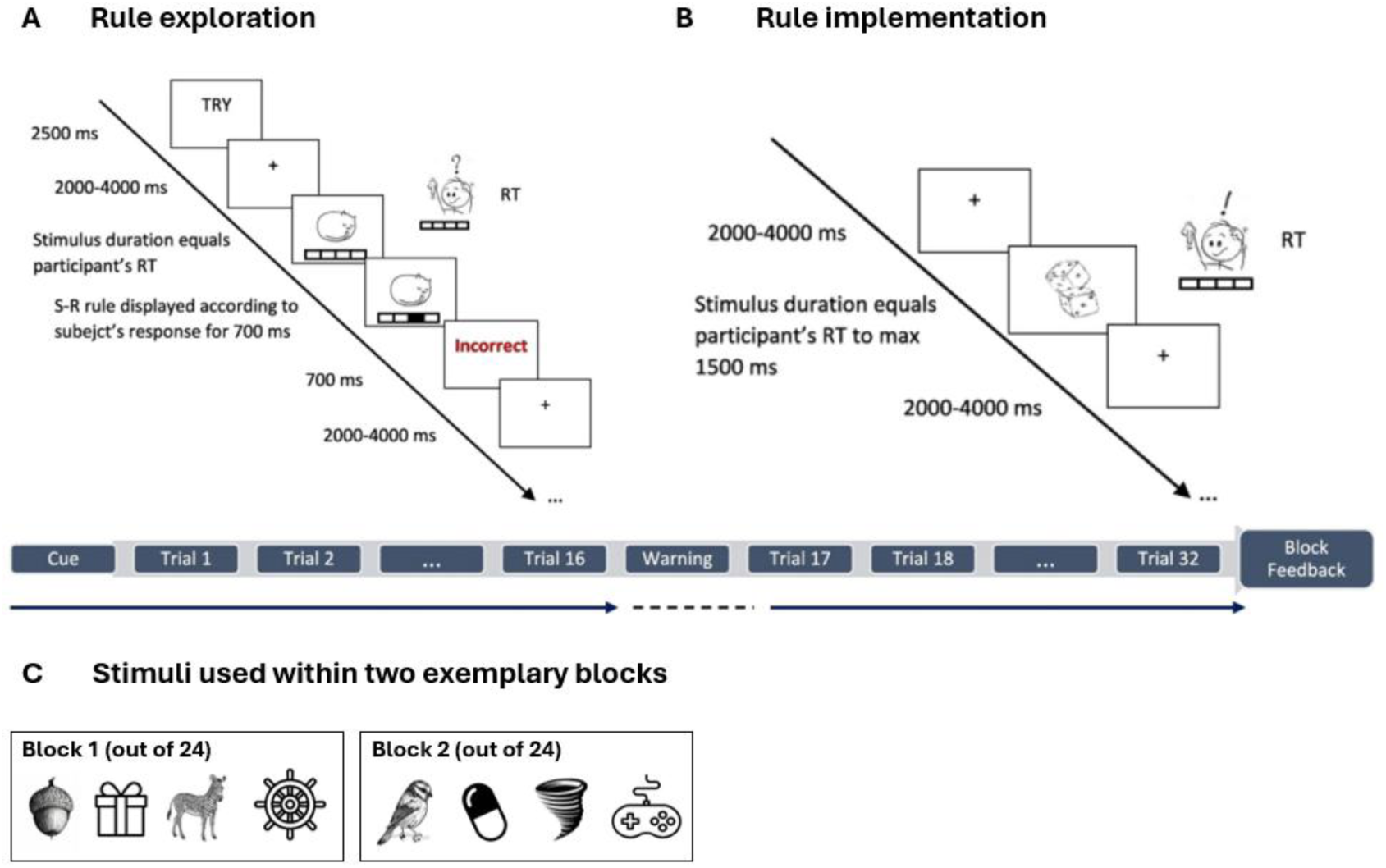
The learning paradigm, displaying the trial-and-error condition. The paradigm featured three conditions in total, including observation based, instruction based, and trial-and-error based learning with 8 learning blocks per condition each involving a novel set of 4 stimuli. However, the present study focused solely on functional connectivity alterations across the trial-and-error based learning condition. (A) Within the rule exploration or learning phase, participants observed the word for ‘TRY’ displayed on the screen, and were then shown a single visual stimulus and four available response keys. Participants were required to press one of the response keys as each stimulus was sequentially displayed. Upon each response, the participants then received response feedback for their correct or incorrect button press. Each of the four stimuli was presented 4 times. (B) Within the rule implementation phase, participants were then presented sequentially with the visual stimuli the same way as before, but now without response feedback. Again, each of the four stimuli was presented 4 times. Figure modified from Fregni *et al*. (2025). (C) Stimuli used within two exemplary learning blocks.

These blocks were randomized across participants, whereby the same run for different participants always consisted of different blocks. This method was utilised in order to ensure that all participants attended the same visual stimuli but in a unique order. A stimulus was presented for the first time as stimulus repetition 1, then a second time as stimulus repetition 2, and so on through all stimulus repetitions. Each block consisted of 32 trials, formed using 4 visual stimuli with 8 stimulus repetitions.

Each experimental block began with a fixation cross displayed on the screen, and after 2.5 seconds, an instruction cue was presented which informed participants on how to respond within the immediately upcoming learning block. These instruction cues consisted of the German words corresponding to “Try!”, “Memorize”, and “Observe”, respectively, which corresponded to the three learning conditions of trial-error, instruction, and observation-based learning. After a variable delay of either 2 or 4 seconds, the first learning trial started. In each trial, the stimuli were presented at the centre of the screen and above the drawing of 4 symbolic response buttons, represented by 4 rectangles placed immediately next to each other.

Within the learning condition that we focused on here, namely the trial-error learning condition, participants were asked to respond to the stimulus of each trial using a button press with their index, middle, ring or little finger of their right hand (Figure 1A). Here, each trial lasted a maximum of 1500 ms or until a response was given. For the first 16 trials (i.e. 4 repetitions of each of the 4 stimuli), each time that a button press was performed by the participant in response to the stimulus, feedback was provided to the participant. This feedback consisted of a symbolic visualization of the executed button press response such that the respective rectangle turned black for 700 ms. This was followed by the German words “Korrekt!” or “Falsch”, depending on whether the feedback was correct or incorrect, respectively. This feedback lasted another 700 ms and was depicted in blue for correct trials and in red for incorrect trials. On occasions when participants failed to submit a button press response within a trial, they would receive a 700-ms red message on the screen warning them to respond. Piloting data indicated that the subject reached asymptotic accuracy at the end of these first 16 trials. This was confirmed in the actual experiment (Figure 3).

For another 16 trials (i.e., 4 more repetitions of each of the 4 stimuli), response feedback was no longer presented, except the absence of a response triggered a 500-ms red message which warned participants that no button press response had been detected, and each trial stayed on the screen until a response was made or for a maximum duration of 1500 ms. These final 16 trials were identical for the three learning conditions. After completion of all 32 trials, each block ended with individual performance-based feedback which displayed the participant’s block accuracy for 3 seconds. This was calculated using the percentage of correct trials within that individual block. Finally, at the end of the experiment, the total accuracy was displayed for 4 seconds.

All trials within the block were interleaved by a fixation cross with a variable duration between 2 and 4 seconds. At the completion of each functional run, participants had a short break whilst fieldmaps were being acquired. Participants were also informed in advance that good performance would give them further opportunity to earn money, whereby a total accuracy of 85% and above within the correct trials throughout the whole experiment would earn them additional money. This additional payment amounted to 1 extra euro for accuracy between 85 to 89%, 2 euros for 90 to 94% accuracy, and 3 extra euros for an overall performance greater than 95% accuracy.

Before the experiment began within the MRI machine, participants completed a practice experiment to ensure understanding of the task. Within this practice session, subjects had to complete one block per learning condition. Across the real experiment, the 12 (4 per practice block) visual stimuli that were utilised within the practice session were not reused.

The experiment was created and performed using E-Prime 3.0 software (Psychology Software Tools, Pittsburgh, PA. (Schneider, 2002). Stimuli sequences and experimental lists were created in MATLAB (version R2020b, 9.13.0 Natick, Massachusetts) and RStudio (R Core Team, 2021).

### MRI Data Acquisition

The MRI data was originally acquired within a Siemens MAGNETOM 3T Trio Tim scanner, using a 32-channel head coil. A BOLD sequence was utilised (TR = 2070 ms, TE = 25 ms, flip angle = 80°, FoV = 192 mm, voxel size 3.0 × 3.0 × 3.2 mm, EPI factor = 64)) which enabled good frontal resolution and minimal ghosting. 36 slices per volume were acquired in a descending sequential order, with each functional acquisition lasting about 25 minutes. This was followed by a GRE EPI fieldmap sequence with the same voxel size as used within the BOLD sequence, for correction of field inhomogeneities of the BOLD acquisition, namely (TR = 439 ms, TE 1 = 5.32 ms, TE 2 = 7.78 ms, flip angle = 45°, FoV = 192 mm, voxel size 3.0 × 3.0 × 3.2 mm). After four functional runs were completed, a high-resolution T1-weighted anatomical image with full brain coverage was then acquired for every subject in the study. This was achieved using MPRAGE, TR = 1900 ms, TE = 2.26 ms, flip angle = 9°, FoV = 256 mm, voxel size = 1.0 × 1.0 × 1.0 mm.

### Data Analysis

#### Preprocessing

Raw MRI data were converted into BIDS 1.2.1 (Gorgolewski *et al*., 2016, RRID:SCR_016124) format before preprocessing. Conversion was implemented via dcm2bids (Boré *et al*., 2023) and Docker (Merkel, 2014). We performed preprocessing using fMRIPrep 22.0.2 (Esteban *et al*., 2018, RRID:SCR_002502; Esteban *et al*., 2019) based on Nipype 1.8.5 (Gorgolewski *et al*., 2011, RRID:SCR_002502; Gorgolewski *et al*., 2018).

#### Preprocessing of B0 inhomogeneity mappings

A total of 4 fieldmaps were available within the input BIDS structure. A *B_0_* nonuniformity map (or *fieldmap*) was estimated from the phase-drift map(s) measure with two consecutive GRE (gradient-recalled echo) acquisitions. The corresponding phase-map(s) were phase-unwrapped with prelude (Jenkinson *et al*., 2012, FSL 6.0.5.1:57b01774).

#### Anatomical data preprocessing

The T1-weighted (T1w) image was corrected for intensity non-uniformity (INU) with N4BiasFieldCorrection (Tustison *et al*., 2010), distributed with ANTs 2.3.3 (Avants *et al*., 2008, RRID:SCR_004757) and used as T1w-reference throughout the workflow. The T1w-reference was then skull-stripped with a *Nipype* implementation of the antsBrainExtraction.sh workflow (from ANTs), using OASIS30ANTs as target template. Brain tissue segmentation of cerebrospinal fluid (CSF), white-matter (WM) and gray-matter (GM) was performed on the brain-extracted T1w using fast (Zhang *et al*., 2001, RRID:SCR_002823; Jenkinson *et al*., 2012, FSL 6.0.5.1:57b01774). Brain surfaces were reconstructed using recon-all in Freesurfer 7.2.0 (Dale *et al*., 1999, RRID:SCR_001847) and the brain mask estimated previously was refined with a custom variation of the method to reconcile ANTs-derived and FreeSurfer-derived segmentations of the cortical gray-matter of Mindboggle (Klein *et al*., 2017, RRID:SCR_002438). Volume-based spatial normalization to one standard space (MNI152NLin2009cAsym) was performed through nonlinear registration with antsRegistration (ANTs 2.3.3), using brain-extracted versions of both T1w reference and the T1w template. The following template was selected for spatial normalization: *ICBM 152 Nonlinear Asymmetrical template version 2009c* (TemplateFlow ID: MNI152NLin2009cAsym, RRID:SCR_008796) (Fonov *et al*., 2009).

#### Functional data preprocessing

For each of the 4 BOLD runs per subject (across all tasks and sessions), the following preprocessing was performed. First, a reference volume and its skull-stripped version were generated using a custom methodology of *fMRIPrep*. Head-motion parameters with respect to the BOLD reference (transformation matrices, and six corresponding rotation and translation parameters) were estimated before any spatiotemporal filtering using mcflirt (Jenkinson *et al*., 2002, FSL, 6.0.5.1:57b01774). The estimated *fieldmap* was then aligned with rigid-registration to the target EPI (echo-planar imaging) reference run. The field coefficients were mapped on to the reference EPI using the transform. BOLD runs were slice-time corrected to 1.01s (0.5 of slice acquisition range 0s-2.02s) using 3dTshift from AFNI (Cox & Hyde, 1997, RRID:SCR_005927). The BOLD reference was then co-registered to the T1w reference using bbregister (FreeSurfer) which implements boundary-based registration (Greve & Fischl, 2009). Co-registration was configured with six degrees of freedom. Six confounding time-series were calculated based on the *preprocessed BOLD*: framewise displacement (FD), DVARS and three region-wise global signals. FD was computed using two formulations, i.e., absolute sum of relative motions (Power *et al*., 2014) and relative root mean square displacement between affines (Jenkinson *et al*., 2002). FD and DVARS were calculated for each functional run, both using their implementations in *Nipype* (following the definitions by Power *et al*. (2014). The three global signals were extracted within the CSF, the WM, and the whole-brain masks. Additionally, a set of physiological regressors were extracted to allow for component-based noise correction (Behzadi *et al*., 2007). Principal components were estimated after high-pass filtering the *preprocessed BOLD* time-series (using a discrete cosine filter with 128s cut-off) for the two *CompCor* variants: temporal (tCompCor) and anatomical (aCompCor). tCompCor components were then calculated from the top 2% variable voxels within the brain mask. For aCompCor, three probabilistic masks (CSF, WM and combined CSF+WM) were generated in anatomical space. The implementation differs from that of Behzadi *et al*. (2007) in that instead of eroding the masks by 2 pixels on BOLD space, a mask of pixels that likely contain a volume fraction of GM is subtracted from the aCompCor masks. This mask is obtained by dilating a GM mask extracted from the FreeSurfer’s *aseg* segmentation, and it ensures components are not extracted from voxels containing a minimal fraction of GM. Finally, these masks were resampled into BOLD space and binarized by thresholding at 0.99 (as in the original implementation). Components were also calculated separately within the WM and CSF masks. For each CompCor decomposition, the *k* components with the largest singular values were retained, such that the retained components’ time series were sufficient to explain 50 percent of variance across the nuisance mask (CSF, WM, combined, or temporal). The remaining components were dropped from consideration.

The head-motion estimates calculated in the correction step were also placed within the corresponding confounds file. The confound time series derived from head motion estimates and global signals were expanded with the inclusion of temporal derivatives and quadratic terms for each (Satterthwaite *et al*., 2013). Frames that exceeded a threshold of 0.5 mm FD or 1.5 standardized DVARS were annotated as motion outliers. Additional nuisance timeseries were calculated by means of principal components analysis of the signal found within a thin band (*crown*) of voxels around the edge of the brain, as proposed by Patriat *et al*. (2017). The BOLD time-series were resampled into standard space, generating a *preprocessed BOLD run in MNI152NLin2009cAsym space*. First, a reference volume and its skull-stripped version were generated using a custom methodology of *fMRIPrep*. The BOLD time-series were resampled onto the following surfaces (FreeSurfer reconstruction nomenclature): *fsnative*. All resamplings can be performed with *a single interpolation step* by composing all the pertinent transformations (i.e. head-motion transform matrices, susceptibility distortion correction when available, and co-registrations to anatomical and output spaces). Gridded (volumetric) resamplings were performed using antsApplyTransforms (ANTs), configured with Lanczos interpolation to minimize the smoothing effects of other kernels (Lanczos, 1964). Non-gridded (surface) resamplings were performed using mri_vol2surf (FreeSurfer).

Many internal operations of *fMRIPrep* use *Nilearn* 0.9.1 (Abraham *et al*., 2014, RRID:SCR_001362), mostly within the functional processing workflow. For more details of the pipeline, see <u>the section</u> <u>corresponding to workflows in *fMRIPrep*’s documentation</u>.

#### Thalamus Segmentation

We used FreeSurfer and the thalamic probabilistic segmentation algorithm by Iglesias *et al*. (2018) which is incorporated into the FreeSurfer software. Utilising a probabilistic thalamus atlas based upon histological data, this algorithm uses Bayesian inference to segment both the left and right thalamic nuclei of individual subjects into an overall total of 47 subregions (see Figure 2 for the results of an exemplary subject). In a first step, we used SynthSeg (Billot *et al*., 2023) to obtain optimized whole-thalamus segmentation. In a second step, the thalamus was segmented into 23 subregions for the left thalamus and 24 subregions for the right thalamus (with increased stiffness of the mesh from 0.05 to 0.25), as the paracentral nucleus on the left side was not available. Here, a subset of 16 subregions per hemisphere were included in our analysis: the anteroventral (AV), central medial (CeM), centromedian (CM), lateral geniculate nucleus (LGN), lateral posterior (LP), medial geniculate nucleus (MGN), mediodorsal lateral parvocellular (MDl), mediodorsal medial magnocellular (MDm), pulvinar anterior (PuA), pulvinar medial (PuM), pulvinar lateral (PuL), pulvinar inferior (PuI), ventral anterior (VA), ventral lateral anterior (VLa), ventral lateral posterior (VLp) nuclei, and ventral posterolateral (VPL) nuclei. However, 7 subregions on the left side and 8 subregions on the right side could not be identified in all our subjects, due to their extremely small volume in the probabilistic atlas. These included the bilateral central lateral (CL), laterodorsal (LD), limitans (suprageniculate) (L-SG), medial ventral (reuniens) (MV-re), parafascicular (Pf), ventral anterior magnocellular (VAmc), ventromedial (VM), and the right paracentral (Pc) nuclei. As a result, these subregions were excluded from our analysis in all subjects.

**Figure 2A-C.**
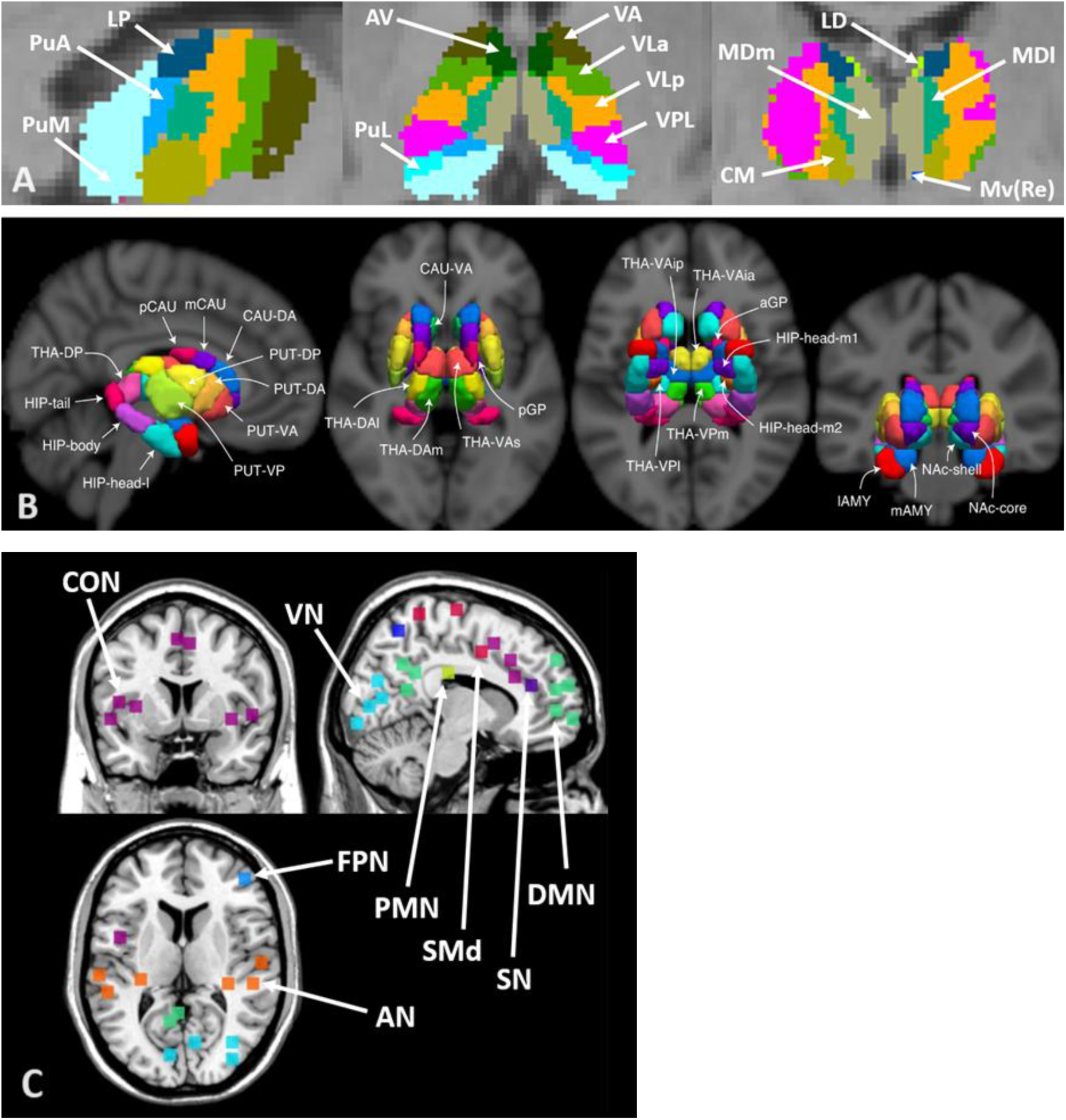
Segmentation of thalamic regions, non-thalamic subcortical regions, and cortical networks. (A) Segmentation of thalamic nuclei using an exemplary subject and a probabilistic atlas by Iglesias *et al*. (2018), displaying 13 thalamic subregions. Subregions include the following: anteroventral (AV), centromedian (CM), lateral dorsal (LD), lateral posterior (LP); mediodorsal lateral (MDl); mediodorsal medial (MDm), nucleus reuniens (Mv(Re)), pulvinar anterior (PuA), pulvinar lateral (PuL), pulvinar medial (PuM); ventral anterior (VA), ventral lateral anterior (VLa); ventral lateral posterior (VLp); and ventral posterolateral (VPL). (B) Subcortical nuclei segmentation of hippocampus, basal ganglia, and thalamus subregions. Figure was adapted from Tian *et al*. (2020). Please note that thalamic subregions were not utilised in our study, as thalamic segmentation was performed using the atlas from Iglesias *et al*. (2018) due to its advantageous segmentation of a larger number of thalamic nuclei and provision of subject-specific segmentation. *Abbreviations:* Hippocampus head medial subdivisions (HIP-head-m1 and HIP-head-m2), hippocampus head lateral subdivision (HIP-head-l), hippocampus body (HIP-body), hippocampus tail (HIP-tail), putamen ventroanterior (PUT-VA), putamen dorsoanterior (PUT-DA), putamen ventroposterior (PUT-VP), putamen dorsoposterior (PUT-DP), caudate ventroanterior (CAU-VA), caudate dorsoanterior (CAU-DA), caudate medial (mCAU), caudate posterior (pCAU), lateral amygdala (lAMY), medial amygdala (mAMY), nucleus accumbens shell (NAc-shell), nucleus accumbens core (NAc-core), globus pallidus posterior (pGP), and globus pallidus anterior (GPa). (C) Cerebral cortex networks obtained from Dworetsky *et al*. (2021). The colours represent the respective networks, namely; Auditory network, AN (orange), cingulo-opercular network, CON (purple); default mode network, DMN (green), fronto-parietal network, FPN (middle blue), parieto-medial network, PMN (yellow), somatomotor dorsal network, SMd (red), salience network, SN (dark blue), and visual network, VN (light blue). These network voxels represent brain regions with highest confidence in network assignment across subjects within the Dworetsky study.

#### Basal Ganglia and Hippocampus Segmentation

Because of their known role in learning, we segmented the putamen, hippocampus, amygdala, caudate nucleus and globus pallidus (Romero *et al*., 2008; Li *et al*., 2011; Kwon *et al*., 2023; Petok *et al*., 2024). To analyse alterations in thalamic connectivity regarding subregions of the hippocampus and components of the basal ganglia, we utilised an atlas created by Tian *et al*. (2020) which segregated 38 bilateral regions (Figure 2B). These ROIs were transformed from MNI space into subject-specific native space using the antsApplyTransforms function based upon the MNI152NLin2009cAsym_to-T1w inverse transformation parameters created during fMRIPrep normalization stage of the pipeline.

#### Cerebral Cortex Networks

We used 12 functional network probability maps that were formed of 153 spherical ROIs, all derived from a recent study by Dworetsky *et al*. (2021). This generated network templates based upon a winner-takes-all procedure across a total of six datasets, available for download at https://github.com/GrattonLab/Dworetsky_etal_ConsensusNetworks. These networks included seven higher-order networks, namely the cingulo-opercular (CO), dorsal attention (DAN), default mode network (DMN), fronto-parietal (FP) network, salience (SN), parietal medial (PMN), and language network (please note that the language network also corresponds to the ventral attention network in other work by Dworetsky *et al*. (2018)) (Figure 2C).

Additionally, further networks were also provided and included in our analysis, namely the visual (VN), somatomotor dorsal (SMd), somatomotor lateral (SMl), and parieto-occipito (PON) networks. These networks were included as sensory and motor networks contribute to goal-directed behaviour through the generation of sensory and motor processes which fuel S-R-O associations (Penalver-Andres *et al*., 2024), whilst the PON generates saccades and has contributions to attention and working memory (Watanabe *et al*., 2010; Galeano Weber *et al*., 2017; Frank *et al*., 2020). Also available from the paper were an additional two networks, including the Temporal Pole (TPole) network and Medial Temporal Lobe (MTL) network. However, the TPole network did not survive correction within the Dworetsky *et al*. (2021) study, and subjects within our own study did not show surviving voxels for the Medial Temporal Lobe (MTL) network. Therefore, neither of these two additional networks were included in our re-analysis.

These cortical network ROIs were subsequently transformed from MNI space into subject-specific native space. This was achieved using the antsApplyTransforms function based on the MNI152NLin2009cAsym_to-T1w inverse transformation parameters, as created during fMRIPrep normalization.

#### Functional data analysis

##### Denoising

After preprocessing, we first performed denoising of the functional data in native space at the single subject level. To this end, we computed a General Linear Model (GLM) for each subject using SPM12 and MATLAB 2021a. The GLM included ‘nuisance’ regressors based on a subset of the confounding variables previously extracted by fMRIPrep. The residual timeseries were saved to disk for further processing. Consistent with the fmridenoise pipleline 24HMPaCompCorSpikeReg, we included the following confounding variables. First, we included the 3 translational and 3 rotational motion parameters and their quadratic terms as well as their derivatives and the quadratic terms of the derivatives. Second, we included the first 5 principal components of the CompCor timeseries within CSF (’c_comp_cor’) and the first 5 principal components of the CompCor timeseries within white matter (’w_comp_cor’). Third, we included the DVARS timeseries and the framewise displacement (‘FD’) timeseries.

##### Modelling task-related activity

In preparation for the functional connectivity analysis based on the beta-series correlation approach, single-trial BOLD activity was estimated. For this we used the Least-Squares-Separate (LSS) modeling approach (Mumford *et al*., 2012; Mumford *et al*., 2014).

For each experimental trial, correct and incorrect trials alike, we ran a separate GLM that included the event of interest, i.e. a single trial and as separate regressors, all the other trials of the same learning condition (e.g., trial-and-error learning), the trials of the other two conditions (observation and instruction-based learning), and the instruction cue, again separately for each learning condition. This amounted to a total of 7 regressors per LSS model. Our regressors were created by convolving stick functions which were synchronized to the stimulus onset using the default canonical HRF within SPM12. The LSS analysis comprised a total of 768 single-trial GLMs (32 trials x 24 blocks) across 4 functional runs and included a high-pass filter at a 200s cut-off. The first 3 scans per run were discarded before starting the analysis.

This analysis was computed with in-house scripts and included functions of the SPM12 software package Statistical Parametric Mapping (RRID:SCR_007037) within MATLAB (version R2020b, The MathWorks Inc. 2020) (RRID:SCR 00162).

##### Calculating functional connectivity

Alterations in functional connectivity were analyzed using the beta-series correlation approach based on the single-trial beta estimates averaged across all voxels within a region of interest (Rissman *et al*., 2004; Abdulrahman & Henson, 2016; Di *et al*., 2021). Previously we focused on the gradual strengthening of habitual responding (which by definition only involves correct responses) over the course of many repetitions (Jarrett *et al*., 2025). Since the goal of the present study was to zoom into the initial phase of this process spanning only the first few repetitions, we again exclusively analyzed correct trials to avoid confounding with neural mechanisms specifically associated with incorrect trials.

Connectivity between two regions was defined as the correlation between the single-trial betas (i.e,. beta-series) for each region. Beta-series correlations were computed for each correct stimulus-repetition separately. To determine the functional connectivity alterations across correct repetitions we computed a linear contrast across correct repetitions 1 to 7. Note that we omitted the final repetition 8 due to the rarity of subjects performing all eight trials correctly.

##### Functional network analysis (ROI-to-ROI connectivity)

In preparation for a ROI-to-ROI functional network analysis regarding learning-related connectivity alterations, we computed the aforementioned linear connectivity contrast for each pair of ROIs for each subject. This resulted in an 81×81 ROI-to-ROI connectivity matrix for each individual subject involving 6480 pairwise connectivity alterations, excluding connectivity of a region with itself. These connectivity matrices were then used as inputs for the group-level functional network analysis (FNC) as implemented in the CONN toolbox (Release 20.a) (Nieto-Castanon, 2020). In a first analysis step, FNC organises the ROIs into networks, as based upon functional connectivity similarity metrics which exist between all ROI-to-ROI pairings, through the use of complete-linkage hierarchical clustering (Sorensen, 1948). In a second step, FNC then analyses all connections between all ROI pairs, for both within- and between-network connectivity sets (Jafri *et al*., 2008). FNC then performs a multivariate parametric GLM analysis for all connections, which generates an F-statistic for each pair of networks with an associated uncorrected cluster-level *p-value*. This is defined as the likelihood of a randomly-selected pair of networks showing equal or greater effects than those observed within the currently considered pair of networks, under the null hypothesis. Finally, a FDR-corrected cluster-level p-value (Benjamini & Hochberg, 1995) is defined as the expected proportion of false discoveries present for all network pairs with similar or larger effects across the total set of pairs (Nieto-Castanon, 2020). We used the default setup with a cluster threshold of p < 0.05 p-*FDR* corrected, alongside a cluster-forming connection threshold of p < 0.05 p-uncorrected.

## Results

### Behavioural Results

As expected, across trial-and-error learning trials, a significant increase regarding accuracy across all eight correct stimulus repetitions was observed (F(3.7,284.8)=1298.2; p<.001; Greenhouse-Geisser corrected) (Figure 3). The steepest accuracy increase occurred from 1^st^ to 3^rd^ stimulus repetitions, before participants’ accuracy values reached ceiling levels around repetition 3 or 4.

**Figure 3.**
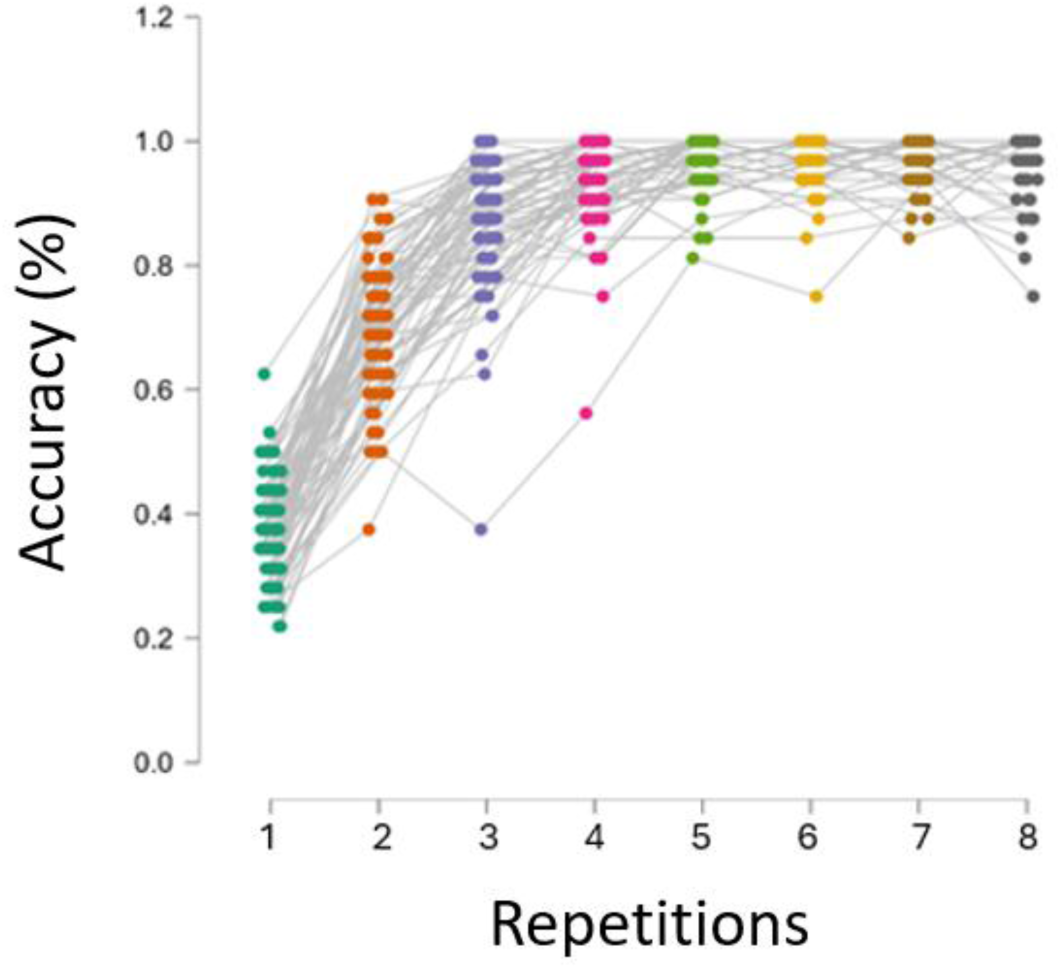
Behavioural results for the trial-and-error learning trials. Accuracy across all eight stimulus repetitions within trial-and-error learning trials. The accuracy ranges from 0 (0% correct responses) to 1 (100% correct responses). Each coloured dot represents the accuracy value for one subject at each repetition. Grey lines connect the dots for each individual subject. The plot was created using JASP (version 0.18.3).

### Learning Related Functional Connectivity Alterations

All results include linear functional connectivity alterations which occurred across correct trial-and-error learning trials from correct repetition 1 to 7. As aforementioned, correct repetition 8 was omitted due to low numbers of participants producing 8 correct responses (typically, errors were made immediately within the first two repetitions). Hence in the majority of blocks they would not reach the maximal number of 8 correct repetitions. Alterations in functional connectivity occurred between numerous higher order thalamic nuclei, cortical networks and putamen subregions (Figure 4). These alterations included increasing functional connectivity between higher order thalamic nuclei with the DMN and salience network, decreasing functional connectivity between higher order thalamic nuclei with FPN, CON and putamen subregions, with these alterations being predominantly centred on mediodorsal nuclei subregions. For visualization purposes, selected alterations are presented individually in Figure 5A-F, and abbreviations are available within Supplementary Table 1. Finally, for detailed statistical information regarding all individual ROI-to-ROI and cluster functional connectivity alterations, please refer to Supplementary Table 2.

**Figure 4.**
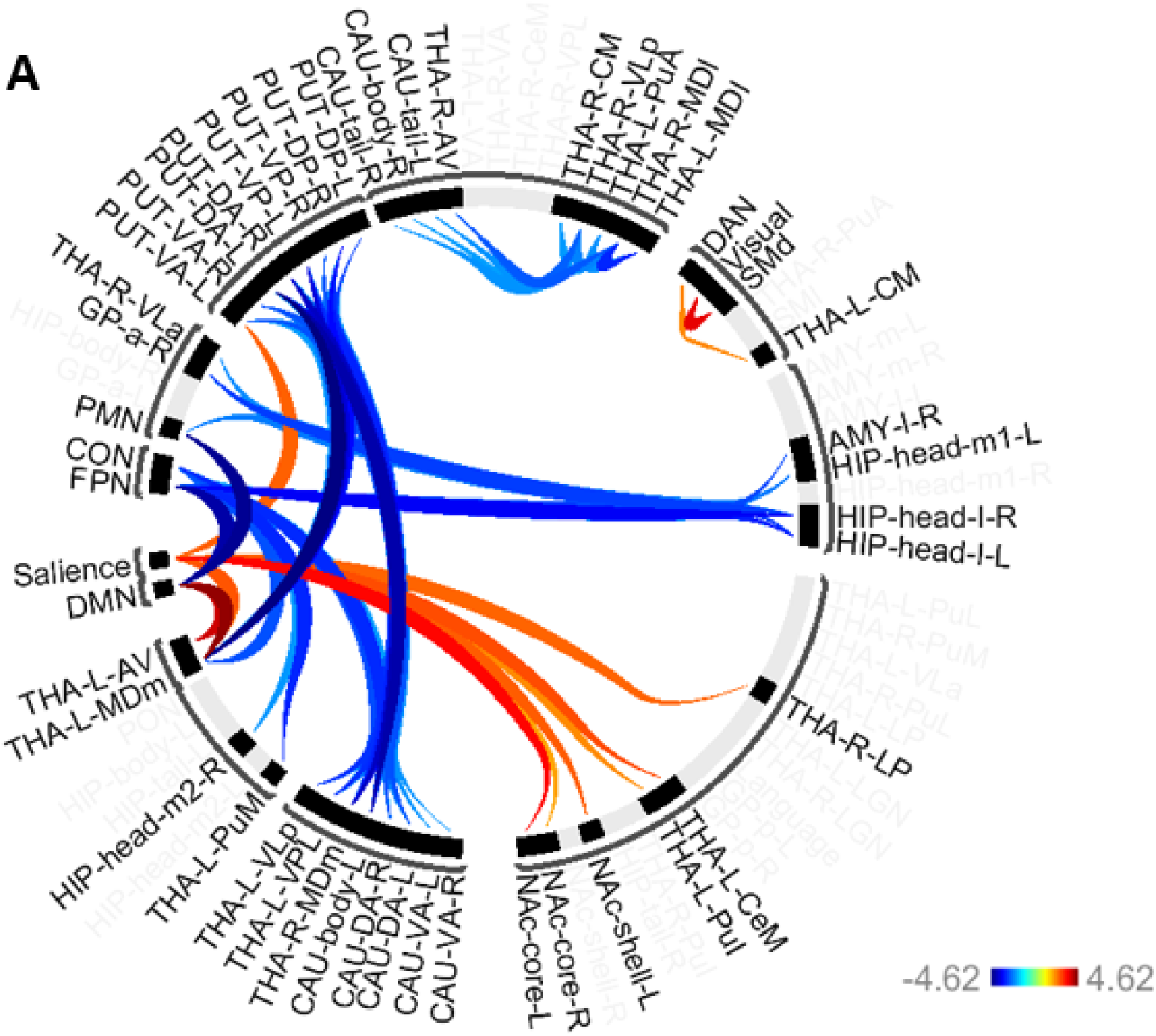
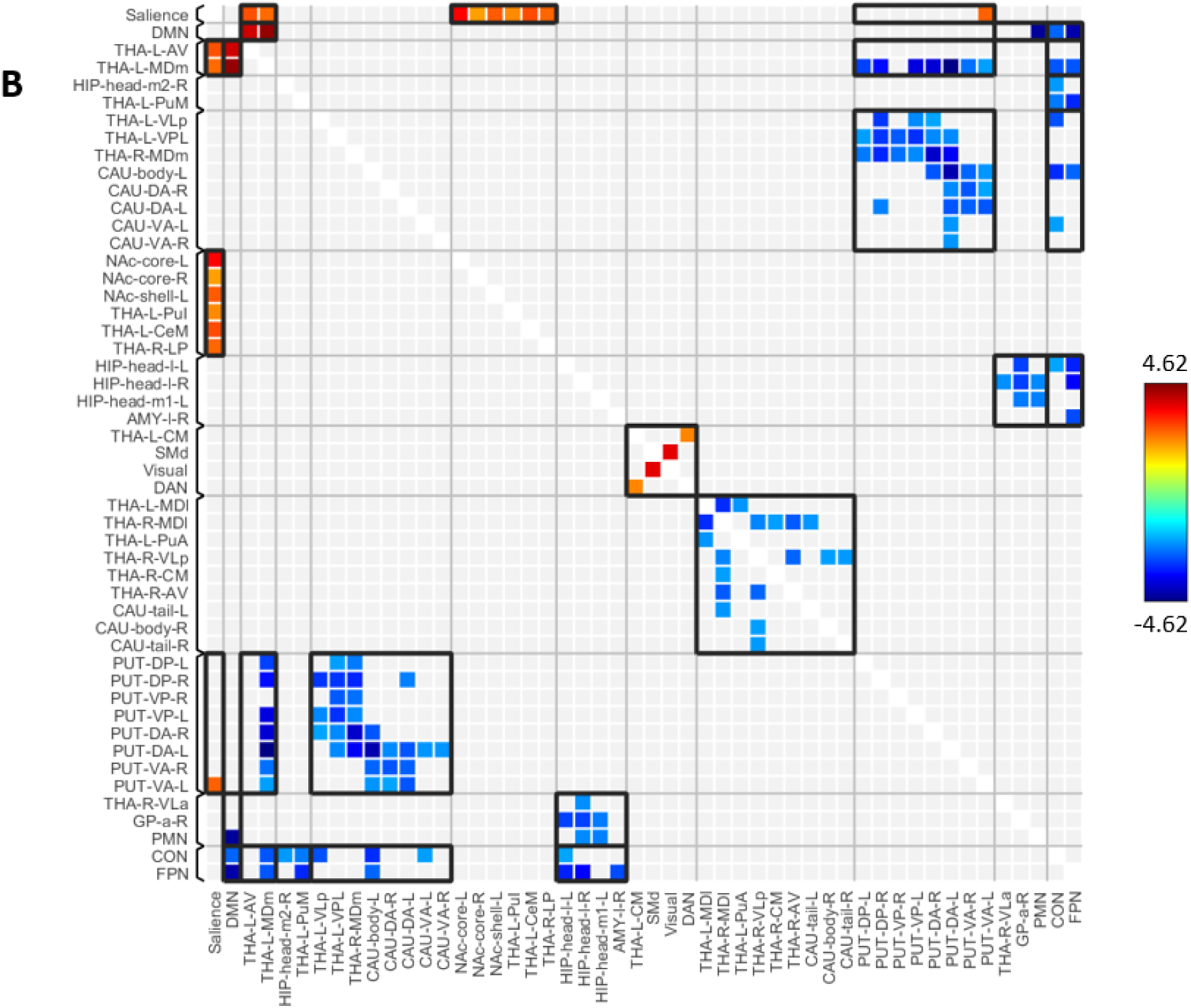
Significant functional connectivity alterations amongst thalamic subregions, subcortical structures, and cortical networks across initial associative S-R learning trials. Colder colours (blue) show decreasing functional connectivity, and darker colours (red) show increasing functional connectivity. The black brackets represent cluster boundaries, as determined by CONN’s hierarchical clustering algorithm. (A) Circular graph representation of the significant connectivity alterations. Darker shaded colours indicate stronger functional connectivity alterations (B) Matrix display representation of the significant connectivity alterations. (*Abbreviations: Organised based on 3 sets of ROIs: (1) THA, thalamus: AV, anteroventral; LGN, lateral geniculate nucleus; LP, lateral posterior; MDl, mediodorsal lateral parvocellular; MDm, mediodorsal medial magnocellular; MGN, medial geniculate nucleus; PuA, pulvinar anterior; PuI, pulvinar inferior; PuL, pulvinar lateral; PuM, pulvinar, medial; VA, ventral anterior thalamus; VLa, ventral lateral anterior; VLp, ventral lateral posterior; VPL, ventral posterolateral. (2) SUBCORTICAL: a, posterior; AMY, amygdala; CAU, caudate; DA, dorsoanterior; DP, dorsoposterior; GP, globus pallidus; HIP, hippocampus; NAc, nucleus accumbens; PUT, putamen; VA, ventroanterior; VP, ventroposterior. (3) CEREBRAL CORTEX: CON, cingulo-opercular; DAN, dorsal attention network; DMN, default mode network; FPN, frontoparietal; p, posterior; PMN, parietal medial network; PON, parieto-occipital network; SMd, somatomotor dorsal; and SMl, somatomotor lateral network. The colour bar refers to the T values across each ROI in the connectivity analysis)*.

**Figure 5A-F.**
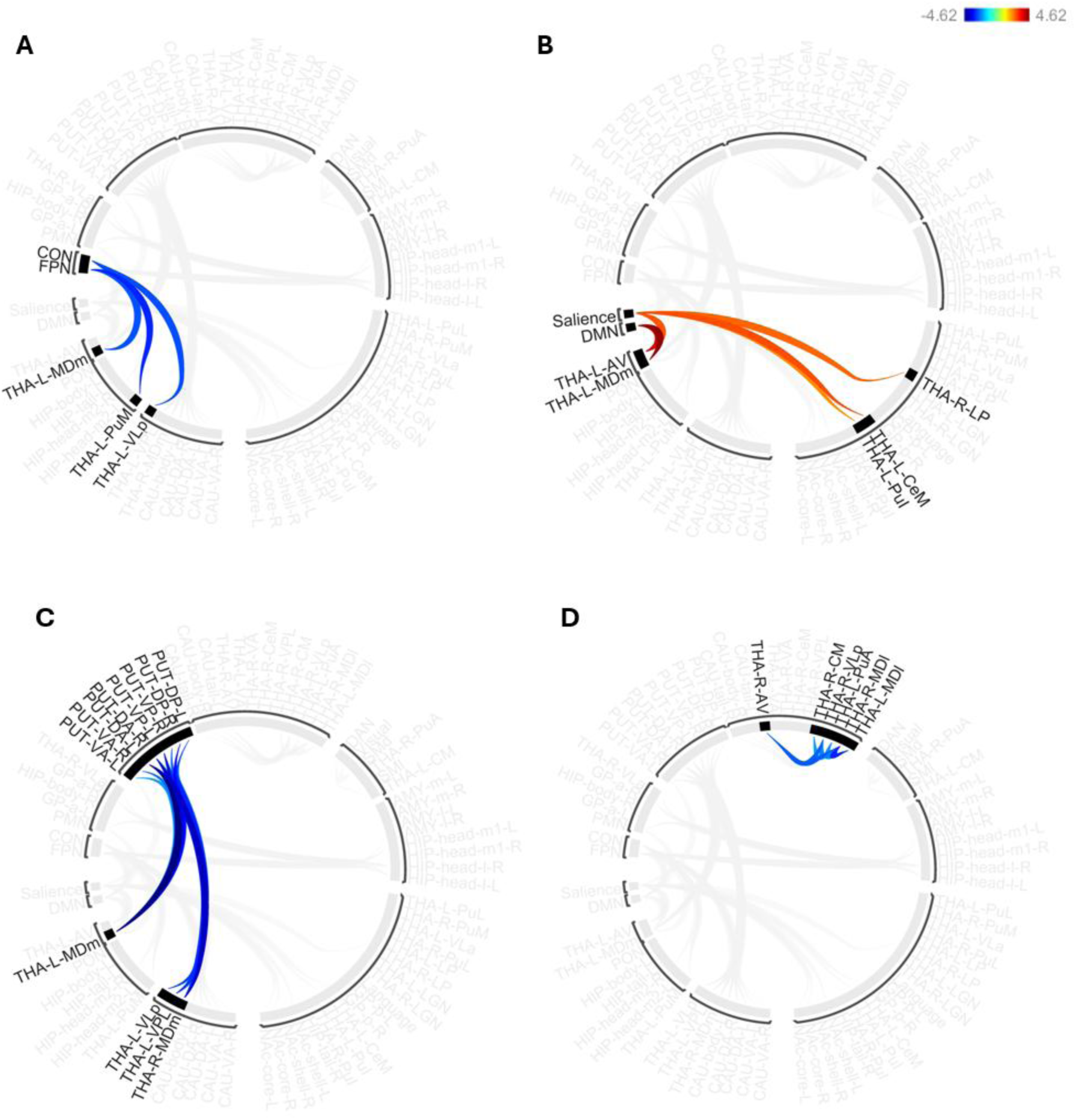
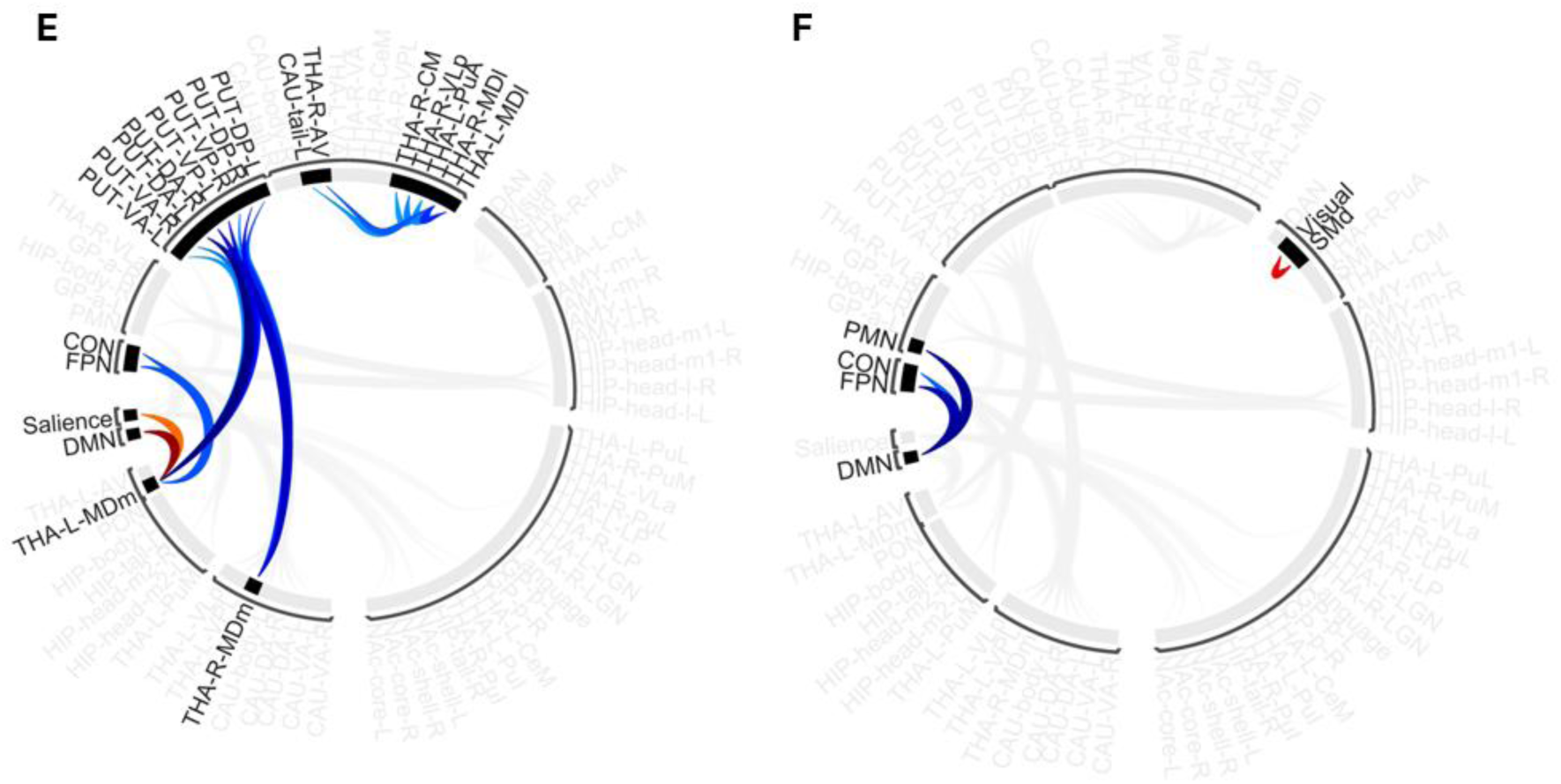
Specific functional connectivity alterations across initial learning trials between thalamic nuclei with non-thalamic subcortical and cortical network regions of interest. The colder colours (blue) show decreasing functional connectivity, and darker colours (red) show increasing functional connectivity. The black brackets represent cluster boundaries, as determined by CONN’s hierarchical clustering algorithm. (A) Decreasing functional connectivity occurred between left higher order thalamic nuclei with both the FPN and CON. (B) Increasing functional connectivity was observed between predominantly left thalamic nuclei with the salience network and DMN. (C) Decreasing functional connectivity was found between predominantly left higher order thalamic nuclei with putamen subregions. (D) Decreasing functional connectivity occurred amongst primarily higher order thalamic nuclei and CSTC nodes. (E) Altogether, these functional connectivity alterations predominantly occurred between mediodorsal thalamic subregions with other networks and regions. (F) Lastly, the DMN exhibited decreasing functional connectivity with task-positive networks whilst sensorimotor networks increased functional connectivity.

### Hierarchical clustering of ROIs (Figure 4A and 4B)

The default hierarchical clustering of ROIs within our analysis showed clustering which was mostly consistent with known regional functions (Figure 4A and 4B; clusters visualized by black brackets). Subcortical subregions generally tended to cluster amongst their own respective subcortical clusters, such that putamen subregions were clustered within a putamen cluster, as were caudate nucleus, nucleus accumbens, and hippocampal subregions. The thalamus, however, showed more complex hierarchical clustering, in which higher order nuclei tended to cluster together but showed more specific patterns of connectivity alterations alongside, or contrasting with, other thalamic nuclei.

Below, the specific connectivity patterns for relevant regions are described in detail as shown in Fig. 5 (A-F).

### Left higher order thalamic nuclei show decreasing functional connectivity with the Frontoparietal Network (FPN) and Cingulo-Opercular Network (CON) across initial learning (Figure 5A)

Higher order thalamic nuclei and CSTC loop components showed decreasing functional connectivity with the FPN and CON during initial learning (Figure 5A). Specifically, these alterations occurred solely amongst left lateralised thalamic nuclei, namely between the FPN with MDm and PuM subregions, and between the CON with MDm, PuM, and VLp subregions.

### Predominantly left higher order thalamic nuclei showed increasing functional connectivity with Default Mode (DMN) and Salience Networks (SN) during initial learning (Figure 5B)

Higher order thalamic nuclei showed increasing functional connectivity with DMN and SN during the initial learning phase (Figure 5B). Specifically, these higher order thalamic nuclei included AV, MD, and pulvinar nuclei (Senatore, 2012; Bennett *et al*., 2019), but also LP and CeM nuclei. The AV and MD nuclei showed increasing functional connectivity with both the DMN and SN, whilst additional increases were also observed between the SN with PuI, LP, and CeM nuclei. Notably, these alterations occurred amongst left lateralised thalamic subregions, with the sole exception of the right LP nucleus.

### Predominantly left thalamic nuclei show decreasing functional connectivity with the putamen across initial learning (Figure 5C)

Thalamic nuclei almost solely on the left side showed decreasing functional connectivity with numerous putamen subregions across initial learning trials (Figure 5C). These thalamic nuclei functional connectivity alterations included left VLp and left VPL nuclei, alongside bilateral MDm nuclei, and therefore included both higher order, CSTC loop nodes, and first order nuclei.

### Higher order thalamic nuclei show decreasing intra-thalamic functional connectivity during initial learning (Figure 5D)

Across both sides of the thalamus, numerous primarily higher order thalamic nuclei or thalamic CSTC nodes exhibited decreasing functional connectivity with each other during initial learning (Figure 5D). These featured alterations across bilateral MDl, left PuA, right VLp, right AV, and right CM. These nuclei included many of the same nuclei which had shown altered functional connectivity with cortical networks and putamen subregions, namely the bilateral MDl, alongside right AV and right VLp (though this had been lateralised to the left side regarding cortical networks and putamen subregions).

### The MD nuclei exhibited major functional connectivity alterations with cortical networks, putamen subregions, and other higher order thalamic nuclei during initial learning (Figure 5E)

The majority of functional connectivity alterations occurred through MD subregions (Figure 5E). These alterations differed based upon MD subregion, as the bilateral MDm exhibited decreasing functional connectivity with putamen subregions, whilst the left MDm additionally showed decreasing functional connectivity with the CON and FPN and increasing functional connectivity with the DMN. Conversely, the bilateral MDl showed decreasing functional connectivity with other thalamic nuclei and the caudate.

### The DMN shows decreasing functional connectivity with task-positive networks, whilst attention and sensorimotor networks show increasing functional connectivity during initial learning (Figure 5F)

The DMN exhibited decreasing functional connectivity with numerous task-positive networks, including FPN, CON, and PMN networks (Figure 5F). In contrast, increasing functional connectivity was observed between the SMd and visual network.

## Discussion

Our results suggest that within the initial phase of trial-and-error learning, diverse thalamic nuclei exhibit dynamic functional connectivity alterations during the early strengthening of the correct stimulus-response (S-R) associations. Firstly, thalamic nuclei exhibited decreasing functional connectivity with the frontoparietal network (FPN), cingulo-opercular network (CON), putamen and other higher order thalamic nuclei, alongside increasing functional connectivity with the default mode network (DMN) and salience network (SN). Furthermore, these thalamocortical, thalamo-putamen and intrathalamic functional connectivity alterations predominantly occurred amongst mediodorsal thalamic subregions, particularly the left mediodorsal medial nucleus (MDm). This suggests that the MD and its cortical and subcortical connections are particularly important for the learning of novel goal-directed behaviour (Halassa & Kastner, 2017; Wolff & Halassa, 2024).

### Left higher order thalamic nuclei show decreasing functional connectivity with frontoparietal and cingulo-opercular networks associated with decreasing cognitive load

Part of the pulvinar is involved in visual processing (and constitutes the “visual thalamus” together with the LGN) (Saalmann & Kastner, 2015), whilst the VL thalamus is involved in motor control (constituting the “motor thalamus”) (Hintzen *et al*., 2018). In contrast, the MD has roles within attention and memory: it serves as a hub for higher-order networks including the DMN and SN, but also executive control networks (Kawabata *et al*., 2021), such as the FPN and CON (Shen *et al*., 2020; Newbold *et al*., 2021), and could be considered to be part of a “cognitive thalamus” (Li *et al*., 2022). In addition, the MD also has an established role as a “limbic thalamus”, in combination with the anterior nuclei regarding its memory processes (Vertes *et al*., 2015). Notably, all of these nuclei showed decreasing functional connectivity with the FPN and CON, and these networks are comprised within the “dual-network” hypothesis – which argues that each network has functionally distinct roles within cognitive control. Here, the CON maintains task rules and monitors performance while the FPN facilitates adjustments to feedback on a trial-to-trial basis (Shen *et al*., 2020). Our results suggest that across learning, there was a decreasing need for communication between these thalamic nuclei and cognitive control networks which reflect decreasing control demands.

In support of this, the FPN shows greater activation during higher cognitive load versus lower cognitive load (Sörqvist *et al*., 2016), and within our current study, cognitive load would likely have decreased overall across the trials for two reasons. Firstly, during earlier learning trials participants needed to explore the space of potential S-R associations and this would have necessitated the holding, monitoring, and updating of numerous potential versus established S-R associations within working memory. Then, as learning progresses, the space of potential S-R associations, and hence, working memory load, decreases. Therefore, on the neural level, this is consistent with the decreasing engagement of FPN and CON along with the diminishing functional connectivity observed between the FPN and higher order thalamic nuclei observed within our study. The decreasing functional connectivity with these selected thalamic nuclei may have been due to their own cognitive, visual and motor functions, suggesting that these nuclei assisted these networks to a greater extent during the earlier stages of higher cognitive load.

### Predominantly left higher order thalamic nuclei show increasing functional connectivity with Default Mode and Salience Networks during initial learning

The pulvinar and LP nuclei support visual processing, the CeM nucleus supports arousal and attention (Schiff et al., 2013; Allen et al., 2016), and both the MD and AV nuclei constitute the limbic thalamus (Wolff et al., 2015) and form the extended limbic system (Grodd et al., 2020). Both MD and anterior nuclei not only play crucial roles within memory and learning (Perry et al., 2021), but they also play important roles in the control of attention and salience (Menon & Uddin, 2010; Zhou et al., 2021), and they connect to numerous cortical networks such as the DMN and SN (Beaty et al., 2021; Kawabata et al., 2021). Both networks are critical for generating goal-directed behaviour and cognitive functions (Andrews-Hanna et al., 2014; Wang et al., 2021; Kesby et al., 2023; Schimmelpfennig et al., 2023), but are known to contribute to these processes in vastly different ways.

Firstly, the DMN plays a crucial role within goal-directed behaviour due to its involvement in internally-generated cognition, including goal-setting and planning (Fingelkurts et al., 2012; Xu et al., 2016; Davey & Harrison, 2018; Vatansever et al., 2018; Dixon et al., 2022). In addition, the DMN supports memory-guided cognitive control (Wen et al., 2020), such as through memory replay (Kaefer et al., 2022). Memory replay is crucial for memory consolidation and preservation (Jafarpour et al., 2017; Wimmer et al., 2020; Wimmer et al., 2023), and within our current study, memory replay is likely to have played a key strategy throughout the learning task. Successful performance across trials required participants to actively memorise the correct S-R associations as they were being discovered and reinforced in memory, and it is therefore also noteworthy that functional connectivity increased between the DMN, MD and anterior thalamic nuclei as these thalamic nuclei are themselves nodes within memory networks and exhibit activity that is modulated by cognitive load (Grodd et al., 2020).

Secondly, the SN also plays a vital role in generating goal-directed behaviour. The SN is a cognitive control network which is critical for identifying the relevance of stimuli and determining the focus of attention (Zhao et al., 2021), serving as a filter for the allocation of attentional resources through top-down modulation (Hegarty et al., 2020), and enabling sustained attention (Steimke et al., 2017). In this way, the SN and its increasing functional connectivity with visual thalamic nuclei may have enabled participants to focus their attention on each stimulus across the trials. The salience of the stimuli may have become stronger with increased stimulus repetitions due to the strengthening of the S-R association and their increasing association with a rewarding outcome. This is also consistent with one of the main roles of the SN as being a network which processes rewards for goal-directed behaviour (von Rhein et al., 2017), alongside that the thalamus is a known node of the salience network – with the MD thalamus forming a loop circuit with the SN; a circuit which has been argued to be central to cognitive control (Peters et al., 2016; Zhou et al., 2021).

### Higher order thalamic nuclei show decreasing functional connectivity with the putamen across initial learning associated with motor selectivity during goal-directed action

The putamen generates and regulates motor movements, and is a critical component of the motor circuit within the basal ganglia (Luo et al., 2019). Together with the caudate nucleus it forms the striatum, which is the primary input structure within the basal ganglia, and the putamen receives mainly excitatory inputs from the cortex, thalamus, and brainstem in the form of potential “action requests” (Buxton et al., 2017; Gerfen, 2022). When activated by the motor cortex, the striatum goes on to activate either direct or indirect pathways (Gerfen, 2022) which generate or inhibit motor movement, respectively. The mechanisms through which this is achieved is via the downstream activation or inhibition of intrinsic and efferent basal ganglia nuclei, and these ganglia nuclei have their own connections with the VL thalamus. This ultimately leads to VL thalamic excitation or inhibition (Rocha et al., 2023), and as the VL nuclei form the motor thalamus (Hintzen et al., 2018), they are reciprocally connected to cortical motor regions (Haber, 2011). These nuclei serve as the central integrator for motor control (Tlamsa & Brumberg, 2010), and form a VA/VL complex with the ventral anterior thalamus (Pelzer et al., 2017) which are themselves nodes within the direct and indirect pathways. Here, they receive innervation from basal ganglia regions and feedback to the motor cortex, generating or inhibiting motor actions, respectively (Young, 2024). When active (achieved through disinhibition), the VL thalamus projects excitatory projections back to the cortex to facilitate movement, thereby constituting the direct pathway within the basal ganglia motor circuit. In contrast, inactivation of the VL thalamus causes motor suppression (Rocha et al., 2023) and constitutes the indirect pathway.

In light of this, and given the decreasing functional connectivity that we observed between the VL thalamus and putamen, this interplay between regions likely influenced the activation and deactivation of direct and indirect pathways within the basal ganglia motor system across initial trial-and-error learning. Successful performance likely involved the incremental exclusion of competing response candidates for each of the stimuli, as each stimulus and its correct response became increasingly strengthened. Specifically, within the earliest repetitions, high competition between alternative response options would have necessitated greater control over responses, such that a single motor response could be selected and executed. This may therefore have been reflected by the strong connectivity that was observed between the thalamus and putamen within earlier repetitions, as consistent with the co-activation of the putamen and VL thalamus within the direct pathway. Then, across later repetitions, this competition would have decreased as a single response to the stimulus would have become dominant, and would be reflected by the weaker connectivity that came to exist between the thalamus and putamen. Overall, this functional connectivity alteration between VL thalamus and putamen therefore likely reflects a changing balance between excitatory and inhibitory motor processes.

In addition to reduced functional connectivity between the putamen and VL thalamus, bilateral MDm nuclei also showed diminishing functional connectivity with almost all putamen subregions. This is interesting as the MD thalamus is a proposed component of frontal cortico-baso-thalamo-cortical (CSTC) loops (Sugiyama et al., 2018) due to its prominent connections with frontal lobe subdivisions (Hummos et al., 2022). Within CSTC loops, the direct and indirect pathways of the basal ganglia system are incorporated (Mathai & Smith, 2011), enabling behavioural responses to be modulated. Accordingly, MD nuclei may play critical roles within the generation of flexible, goal-directed behaviour through outcome-driven responding (Morceau et al., 2022), via the modulation of putamen activity.

### Comparison with a previous study which investigated the transition from goal-directed behaviour toward more habitual behaviour within trial-and-error associative learning

Previously, we had performed a similar study to the current study (Jarrett et al., 2025), which also investigated associative visual stimulus-response learning. That paradigm was designed to generate a transition from goal-directed behaviour toward more automatic (or habitual) behaviour, owing to its use of 80 repetitions per stimulus instead of the mere eight stimulus repetitions of the current study. This previous study therefore contrasts with the present study which investigated the early phase of S-R learning supposedly dominated by goal-directed processes. In Jarrett et al. (2025), we had also investigated functional connectivity alterations between thalamic nuclei, non-thalamic subcortical regions, and cortical networks across correct trials using fMRI and observed numerous alterations in functional connectivity, including: (1) decreasing FC between the frontoparietal network and higher order thalamic nuclei; (2) increasing FC between the cingulo-opercular network and pulvinar nuclei; (3) decreasing FC between the default mode network (DMN) and right mediodorsal nuclei; (4) increasing FC between the DMN and left mediodorsal nuclei; (5) changes in functional connectivity between thalamic nuclei and putamen subregions, and (6) increasing intrathalamic FC.

When contrasted with the results of the Jarrett et al. (2025) study, our present results reveal both common and distinct functional connectivity dynamics during the early phase of S-R learning. Similarities with the current study include decreasing functional connectivity between thalamic nuclei and the FPN, increasing functional connectivity with selected cortical networks, alongside altered functional connectivity between thalamic nuclei and the putamen – however, the specific alterations differed across the studies. For example, in the current study we found increasing functional connectivity between thalamic nuclei, DMN and salience networks, whilst for learning with more repetitions, we found increasing functional connectivity between visual thalamic nuclei and the CON (Cortes *et al*., 2024; Jarrett *et al*., 2025). This suggests that distinct thalamocortical functional connectivity patterns underlie initial versus later learning processes, which is supported by previous research which reports timescale-specific activation of cortical networks (Luu *et al*., 2014; Sorrentino *et al*., 2023; van Es *et al*., 2025). Specifically, these results also suggest that initial learning and goal-directed behaviour are supported by thalamic nuclei and cortical networks which support attention and internal, conscious cognition respectively (Andrews-Hanna *et al*., 2010; de Bourbon-Teles *et al*., 2014; Bodien *et al*., 2019), whilst later learning processes and increasingly automatic behaviour are supported by visual thalamic nuclei and cortical networks which support the implementation of learned task sets (Marek & Dosenbach, 2018). Furthermore, whilst the current study also found decreasing functional connectivity between MD nuclei and all putamen subregions, the Jarrett *et al*. (2025) study found increasing functional connectivity between thalamic nuclei and the left ventroanterior putamen. Here, whilst both the anterior and posterior putamen are both involved in sensorimotor processes, it is the anterior putamen which is critical for the generation of newly acquired habits (Guida *et al*., 2022). Therefore, these findings suggest that initial versus later learning processes are also supported by progressive thalamo-putamen functional connectivity alterations which promote different sensorimotor behaviours. Altogether, these distinct thalamocortical and thalamo-putamen patterns appear to drive goal-directed versus increasingly automatic behaviours within the brain.

## Conclusion

We found that during the initial phase of associative S-R learning, numerous higher order thalamic nuclei exhibited functional connectivity alterations with cortical networks, putamen subregions, and other thalamic nuclei. These alterations are consistent with known functions of thalamic nuclei, cortical networks, and the putamen, and appear to contribute to the generation of goal-directed behaviour during initial trial-and-error learning processes. Here, MD thalamic nuclei appeared to play a particularly prominent role through their altered functional connectivity across cortical networks and putamen subregions.

