## Supplementary material for "Rapid connectivity alterations of thalamic nuclei during initial learning of goal-directed behaviour": N/A

Supplementary Materials

**Supplementary Table 1.** List of subregions and corresponding abbreviations.

| **Regions of Interest and Abbreviations** | | | |
|---|---|---|---|
| **Thalamus** | **Abbreviation** | **Amygdala** | **Abbreviation** |
| Anteroventral | AV | Lateral amygdala | lAMY |
| Central medial | CeM | Medial amygdala | mAMY |
| Centromedian | CM | **Caudate** | |
| Lateral geniculate nucleus | LGN | Ventroanterior caudate | VA CAU |
| Lateral posterior | LP | Dorsoanterior caudate | DA CA |
| Mediodorsal lateral parvocellular | MDl | **Globus Pallidus** | |
| Mediodorsal medial magnocellular | MDm | Anterior globus pallidus | aGP |
| Medial geniculate nucleus | MGN | Posterior globus pallidus | pGP |
| Pulvinar anterior | PuA | **Hippocampus** | |
| Pulvinar inferior | PUI | Hippocampus head, medial division, subdivision 1 | HC head, medial 1 |
| Pulvinar lateral | PuL | Hippocampus head, medial division, subdivision 2 | HC head, medial 2 |
| Pulvinar medial | PuM | Hippocampus head, lateral division | HC head, lateral |
| Ventral anterior | VA | Hippocampus body | HC body |
| Ventral lateral anterior | VLa | Hippocampus tail | HC tail |
| Ventral lateral posterior | VLp | **Nucleus Accumbens** | |
| Ventral posterolateral | VPL | Nucleus accumbens, shell | NAcc, shell |
| **Cortical Networks** | | Nucleus accumbens, core | NAcc, core |
| Cingulo-opercular network | CON | **Putamen** | |
| Default mode network | DMN | Ventroanterior putamen | VA PUT |
| Dorsal attention network | DAN | Dorsoanterior putamen | DA PUT |
| Frontoparietal network | FPN | Ventroposterior putamen | VP PUT |
| Language network | Language | Dorsoposterior putamen | DP PUT |
| Parietal medial network | PMN | | |
| Parieto-occipital network | PON | | |
| Salience network | Salience | | |

| | |
|---|---|
| Somatomotor dorsal network | SMd |
| Somatomotor lateral network | SMl |
| Visual network | Visual |

**Supplementary Table 2.** Detailed results for the group-level functional connectivity analysis, which were obtained using the Functional Network Connectivity multivariate parametric statistics implemented in the CONN toolbox. This table lists significant connectivity clusters for network pairs with $p < 0.05$ *FDR*-corrected across clusters (in bold font) together with univariate statistics for individual connections within each cluster surviving an uncorrected threshold of $p < 0.05$.

| **Analysis Unit** | **Statistic** | **p-unc** | **p-FDR** |
|---|---|---|---|
| **Cluster 1/76** | **F(2,76) = 13.31** | 0.000011 | 0.000844 |
| Connection DMN – Thal-L-MDm | T(77) = 4.44 | 0.000030 | 0.001215 |
| Connection DMN – Thal-L-AV | T(77) = 3.89 | 0.000213 | 0.004157 |
| **Cluster 2/76** | **F(2,76) = 10.65** | **0.000083** | **0.003170** |
| Connection DMN – FPN | T(77) = -4.27 | 0.000055 | 0.001423 |
| Connection DMN – CON | T(77) = -2.58 | 0.011676 | 0.091074 |
| **Cluster 3/76** | **F(3,75) = 6.93** | **0.000351** | **0.008886** |
| Connection DMN – PMN | T(77) = -4.43 | 0.000031 | 0.001215 |
| **Cluster 4/76** | **F(3,75) = 5.79** | **0.001291** | **0.020627** |
| Connection Thal-L-MDm – PUT-DA-L | T(77) = -4.62 | 0.000015 | 0.001152 |
| Connection Thal-L-MDm – PUT-DA-R | T(77) = -3.90 | 0.000203 | 0.005287 |
| Connection Thal-L-MDm – PUT-VP-L | T(77) = -3.80 | 0.000287 | 0.005596 |
| Connection Thal-L-MDm – PUT-DP-R | T(77) = -3.42 | 0.000990 | 0.012867 |
| Connection Thal-L-MDm – PUT-DP-L | T(77) = -2.94 | 0.004310 | 0.048031 |
| Connection Thal-L-MDm – PUT-VA-R | T(77) = -2.48 | 0.015494 | 0.092966 |
| Connection Thal-L-MDm – PUT-VA-L | T(77) = -2.06 | 0.042865 | 0.208966 |
| **Cluster 5/76** | **F(3,75) = 5.73** | **0.001379** | **0.020627** |
| Connection HIP-head-l-R – aGP-R | T(77) = -2.94 | 0.004329 | 0.168848 |
| Connection HIP-head-l-L – aGP-R | T(77) = -2.92 | 0.004643 | 0.181063 |
| Connection HIP-head-l-R – PMN | T(77) = -2.25 | 0.027039 | 0.491814 |
| Connection HIP-head-l-R – Thal-R-VLa | T(77) = -2.19 | 0.031527 | 0.491814 |
| Connection HIP-head-m1-L – aGP-R | T(77) = -2.38 | 0.019784 | 0.509130 |
| Connection HIP-head-m1-L – PMN | T(77) = -2.34 | 0.021980 | 0.509130 |
| **Cluster 6/76** | **F(3,75) = 5.59** | **0.001628** | **0.020627** |
| Connection Thal-L-MDm – CON | T(77) = -2.74 | 0.007550 | 0.061663 |
| Connection Thal-L-MDm – FPN | T(77) = -2.73 | 0.007905 | 0.061663 |
| **Cluster 7/76** | **F(3,75) = 5.34** | **0.002174** | **0.020874** |
| Connection CAU-body-L – PUT-DA-L | T(77) = -4.22 | 0.000067 | 0.005194 |
| Connection Thal-R-MDm – PUT-DA-R | T(77) = -4.01 | 0.000141 | 0.010964 |
| Connection Thal-R-MDm – PUT-DA-L | T(77) = -3.59 | 0.000573 | 0.022362 |
| Connection Thal-R-MDm – PUT-DP-R | T(77) = -3.18 | 0.002094 | 0.054441 |
| Connection CAU-body-L – PUT-DA-R | T(77) = -2.72 | 0.008084 | 0.091258 |

| | | | |
|---|---|---|---|
| Connection CAU-body-L – PUT-VA-R | T(77) = -2.58 | 0.011700 | 0.091258 |
| Connection CAU-DA-L – PUT-VA-L | T(77) = -2.75 | 0.007432 | 0.122868 |
| Connection CAU-DA-L – PUT-DA-L | T(77) = -2.73 | 0.007876 | 0.122868 |
| Connection CAU-DA-L – PUT-VA-R | T(77) = -2.66 | 0.009548 | 0.124126 |
| Connection Thal-L-VPL – PUT-VP-L | T(77) = -3.08 | 0.002896 | 0.134977 |
| Connection Thal-L-VPL – PUT-VP-L | T(77) = -3.08 | 0.002896 | 0.134977 |
| Connection Thal-L-VPL – PUT-DP-R | T(77) = -3.02 | 0.003461 | 0.134977 |
| Connection Thal-R-MDm – PUT-VP-R | T(77) = -2.50 | 0.014604 | 0.162731 |
| Connection Thal-R-MDm – PUT-DP-L | T(77) = -2.41 | 0.018263 | 0.178068 |
| Connection Thal-L-VLp – PUT-DP-R | T(77) = -3.05 | 0.003165 | 0.187225 |
| Connection CAU-body-L – PUT-VA-L | T(77) = -2.16 | 0.034259 | 0.190871 |
| Connection Thal-R-MDm – PUT-VP-L | T(77) = -2.27 | 0.026198 | 0.205400 |
| Connection CAU-DA-R – PUT-VA-R | T(77) = -2.75 | 0.007356 | 0.209436 |
| Connection Thal-L-VPL – PUT-VP-R | T(77) = -2.70 | 0.008598 | 0.223558 |
| Connection CAU-DA-L – PUT-DP-R | T(77) = -2.33 | 0.022656 | 0.224627 |
| Connection Thal-L-VPL – PUT-DA-L | T(77) = -2.25 | 0.027130 | 0.275176 |
| Connection Thal-L-VPL – PUT-DA-R | T(77) = -2.24 | 0.028223 | 0.275176 |
| Connection CAU-DA-R – PUT-DA-L | T(77) = -2.26 | 0.026645 | 0.283437 |
| Connection CAU-DA-R – PUT-VA-L | T(77) = -2.02 | 0.046571 | 0.302711 |
| Connection CAU-VA-R – PUT-DA-L | T(77) = -2.15 | 0.035049 | 0.306211 |
| Connection Thal-L-VPL – PUT-DP-L | T(77) = -2.06 | 0.042300 | 0.366598 |
| Connection Thal-L-VLp – PUT-VP-L | T(77) = -2.25 | 0.027380 | 0.427131 |
| Connection Thal-L-VLp – PUT-DA-R | T(77) = -2.02 | 0.046732 | 0.455640 |
| Connection CAU-VA-L – PUT-DA-L | T(77) = -2.15 | 0.034797 | 0.492267 |
| **Cluster 8/76** | **F(3,75) = 5.33** | **0.002197** | **0.020874** |
| Connection SAN – NAc-core-L | T(77) = 3.45 | 0.000919 | 0.065573 |
| Connection SAN – Thal-L-CeM | T(77) = 2.77 | 0.006977 | 0.121803 |
| Connection SAN – NAc-shell-L | T(77) = 2.63 | 0.010334 | 0.121803 |
| Connection SAN – Thal-R-LP | T(77) = 2.55 | 0.012835 | 0.125144 |
| Connection SAN – Thal-L-Pul | T(77) = 2.20 | 0.031175 | 0.213830 |
| Connection SAN – NAc-core-R | T(77) = 2.00 | 0.049167 | 0.273928 |
| **Cluster 9/76** | **F(3,75) = 4.99** | **0.003295** | **0.026880** |
| Connection SAN – PUT-VA-L | T(77) = 2.61 | 0.010931 | 0.121803 |
| **Cluster 10/76** | **F(3,75) = 4.93** | **0.003537** | **0.026880** |
| Connection SMd – Visual | T(77) = 3.70 | 0.000406 | 0.022911 |
| Connection Thal-L-CM – DAN | T(77) = 2.25 | 0.027264 | 0.471147 |
| **Cluster 11/76** | **F(3,75) = 4.79** | **0.004181** | **0.028889** |
| Connection CAU-body-L – CON | T(77) = -3.10 | 0.002696 | 0.070095 |
| Connection CAU-body-L – FPN | T(77) = -2.60 | 0.011186 | 0.091258 |
| Connection Thal-L-VLp – CON | T(77) = -2.80 | 0.006537 | 0.187225 |
| Connection CAU-VA-L – CON | T(77) = -2.05 | 0.044073 | 0.492267 |
| **Cluster 12/76** | **F(3,75) = 4.46** | **0.006159** | **0.039010** |
| Connection HIP-head-l-R – FPN | T(77) = -3.53 | 0.000708 | 0.055254 |
| Connection HIP-head-l-L – FPN | T(77) = -3.22 | 0.001855 | 0.144727 |
| Connection HIP-head-l-L – CON | T(77) = -2.01 | 0.048477 | 0.479828 |
| Connection lAMY-R – FPN | T(77) = -2.81 | 0.006244 | 0.487064 |

| | | | |
|---|---|---|---|
| **Cluster 13/76** | **F(3,75) = 4.34** | **0.007087** | **0.039892** |
| Connection Thal-L-MDl – Thal-R-MDl | T(77) = -3.14 | 0.002408 | 0.093918 |
| Connection Thal-R-MDl – Thal-R-AV | T(77) = -2.67 | 0.009151 | 0.009151 |
| Connection Thal-L-MDl – Thal-L-PuA | T(77) = -2.15 | 0.034434 | 0.278393 |
| Connection Thal-R-MDl – Thal-R-VLp | T(77) = -2.27 | 0.025728 | 0.335822 |
| Connection Thal-R-MDl – CAU-tail-L | T(77) = -2.13 | 0.036432 | 0.335822 |
| Connection Thal-R-MDl – Thal-R-CM | T(77) = -2.06 | 0.043054 | 0.335822 |
| Connection Thal-R-VLp – Thal-R-AV | T(77) = -2.61 | 0.010945 | 0.409033 |
| Connection Thal-R-VLp – CAU-tail-R | T(77) = -2.12 | 0.036998 | 0.409033 |
| Connection Thal-R-VLp – CAU-body-R | T(77) = -2.03 | 0.045477 | 0.409033 |
| **Cluster 14/76** | **F(2,76) = 5.22** | **0.007496** | **0.039892** |
| Connection SAN – Thal-L-AV | T(77) = 2.71 | 0.008290 | 0.121803 |
| Connection SAN – Thal-L-MDm | T(77) = 2.48 | 0.015481 | 0.134167 |
| **Cluster 15/76** | **F(3,75) = 4.25** | **0.007874** | **0.039892** |
| Connection Thal-L-PuM – FPN | T(77) = -3.16 | 0.002284 | 0.089082 |
| Connection Thal-L-PuM – CON | T(77) = -2.37 | 0.020058 | 0.260753 |
| Connection HIP-head-m2-R – CON | T(77) = -2.10 | 0.038707 | 0.835850 |
